# Standing balance assessment using a head-mounted wearable device

**DOI:** 10.1101/149831

**Authors:** Joseph P. Salisbury, Neha U. Keshav, Anthony D. Sossong, Ned T. Sahin

## Abstract

**Background:** The presence of accelerometers in smartphones has enabled low-cost balance assessment. Smartglasses, which contain an accelerometer similar to that of smartphones, could provide a safe and engaging platform for virtual and augmented reality balance rehabilitation; however, the validity of head-mounted measurement of balance using smartglasses has not been investigated.

**Objective:** To perform preliminary validation of a smartglasses-based balance accelerometry measure (BAM) compared with previously validated waist-based BAM.

**Methods:** 42 healthy individuals (26 male, 16 female; mean age ± SD = 23.8 ± 5.2 years) participated in the study. Following the BAM protocol, each subject performed two trials of six balance stances while accelerometer and gyroscope data were recorded from smartglasses (Google Glass). Test-retest reliability and correlation were determined relative to waist-based BAM as used in the NIH Standing Balance Toolbox.

**Results:** Balance measurements obtained using a head-mounted wearable were highly correlated with those obtained through a waist-mounted accelerometer (Spearman’s rank correlation coefficient = 0.85). Test-retest reliability was high (ICC = 0.85, 95% CI 0.81-0.88), and in good agreement with waist balance measurements (ICC = 0.84, 95% CI 0.80-0.88). Taking into account the total NPL magnitude improved inter-device correlation (0.90) while maintaining test-retest reliability (0.87, 95% CI 0.83-0.90). All subjects successfully completed the study, demonstrating the feasibility of using a head-mounted wearable to assess balance in a healthy population.

**Conclusion:** Balance measurements derived from the smartglasses-based accelerometer were consistent with those obtained using a waist-mounted accelerometer. Given this and the potential for smartglasses in vestibular rehabilitation, the continued development and validation of balance assessment measurements obtained via smartglasses is warranted. This research was funded in part by Department of Defense/Defense Health Program (#W81XWH-14-C-0007, SBIR Phase II contract awarded to TIAX, LLC).

## Introduction

Up to 75% of people over the age of 70 years old experience abnormal postural balance [1], which severely impacts their quality of life and contributes to decline in overall health [2]. Balance assessment can be used to prevent fall-related injury in the elderly [3], as well as inform critical clinical decisions regarding a variety of neurological impairments and movement disorders, including traumatic brain injury (TBI) [4], stroke [5], multiple sclerosis [6], Parkinson’s disease [7], and arthritis [8]. A variety of subjective and objective assessments exist to both identify and characterize balance deficits [9]. Objective standing balance assessments generate valid and reliable quantitative measures through devices such as force platforms, strain gauges, and accelerometers. Accelerometer-based assessments have garnered increased attention due to their widespread availability as a component of consumer smartphones [10]. These smartphone devices routinely contain 9-axis inertial measurement units (IMUs) that include a 3-axis accelerometer, along with a gyroscope and magnetometer. The Balance Accelerometer Measure (BAM) of the NIH Toolbox was developed by the National Institutes of Health (NIH) in order to provide one such low-cost assessment [11], which can now be administered through the use of an iOS (Apple Inc., Cupertino, CA) app. Likewise, the Sway balance application [12] for iOS has gained FDA clearance to assess sway as an indicator of balance using proprietary measures based upon accelerometer measurements obtained during traditional clinical balance stances.

In addition to smartphones, a growing landscape of consumer wearable devices include IMUs with accelerometers. Smartglasses, such as Glass (Google, Mountain View, CA), combine components found in smartphones such as IMUs with a head-mounted display (HMD), camera, microphone, and audio output. Smartglasses could enable self-administered balance assessments and rehabilitative feedback by providing integrated instruction to the user through the HMD while monitoring balance via the IMU.

Despite the potential for smartglasses to provide integrated measurement and training, little research has been conducted regarding IMU-based balance measurement on smartglasses. In this report, we examined the correspondence between head-mounted accelerometer measurements obtained on smartglasses with those obtained on a consumer smartphone device attached to the waist, such as when administering the NIH Toolbox Standing Balance Test.

## Methods

### Subjects

42 healthy individuals participated (**Table 1**). Subjects were recruited from the public and required to be between the ages of 18-39 years old, weigh no more than 250 pounds, and possess normal hearing and normal or corrected-to-normal vision. All participants were free from any preexisting condition that may have altered their ability to balance normally, including multiple sclerosis, Parkinson’s disease, Huntington’s disease, other movement disorders, stroke, cervical spine or physical mobility issues, more than one fall in the past 5 months not as a result of an accident, current pregnancy, dizziness or vertigo, any lower extremity injury that required medical attention in the last three months, any surgeries within the last year, or any medication or other substance that would affect balance. Individuals were also screened for history of a diagnosed seizure disorder (or any seizures within the last 3 years), as well as extreme sensory sensitivity. All participants attested to having no diagnosed macular degeneration, glaucoma, or cataracts, or any chronic or acute conditions resulting in pain, including diabetes or a history of joint replacement. Procedures were approved by Asentral, Inc. Institutional Review Board (Newburyport, MA, USA) and the U.S. Army Human Research Protection Office. Informed consent was obtained from all subjects prior to participation.

**Table 1.** Subject demographics (*n*=42)

|  |  |
| --- | --- |
| Age, years: mean (SD, range) | 23.8 (5.2, 18-37) |
| Gender (Female, percent) | 16 (38) |
| Height, inches: mean (SD, range) | 68 (3, 62-76) |
| Weight, lbs.: mean (SD, range) | 152 (32, 110-241) |

### Experimental setup

Prior to administering the BAM protocol, subjects were outfitted with a gait belt. An Android smartphone (Samsung Galaxy S5, Samsung Galaxy S6, or LG Electronics/Google Nexus 5) was attached to the gait belt using a protective case with clip. The smartphone was attached upright, with the screen was facing away from the subject. The subject was given a pair of Google Glass by the facilitator to also wear. Subjects who normally wore glasses were given the option to wear Google Glass without their regular glasses, or wear Google Glass over their glasses. Subjects were asked to read a sentence on the display screen to confirm the screen was adjusted properly. A test exercise was administered on Glass to ensure subject could: (1) Operate Glass by tapping on the side, and (2) Could hear a tone played from Glass. The BAM protocol was administered as previously described [13]. Briefly, the BAM protocol includes six standing conditions: (1) Solid surface, feet together, eyes open, and (2) eyes closed; (3) Foam surface (Airex Balance Pad, Speciality Foams, Switzerland), feet together, eyes open, and (4) eyes closed; (5) Solid surface, tandem standing, eyes open, and (6) eyes closed (**Fig. 1**). During each stance, all subjects were asked to stand quietly for 60 seconds and to look (in eyes-open conditions) at a symbol placed centrally at eye level one meter from the subject. Subjects were instructed by facilitator regarding stance following the instructions adapted from the NIH Toolbox Standing Balance Test [11, 14]. Stance was also described on the smartglasses display screen. Subjects initiated each set of data collection by tapping the side of the smartglasses. A timer was displayed on Glass showing time remaining and a tone was played at the end of each timed stance. All subjects completed two trials of all stances on the same day.

**Fig. 1.**
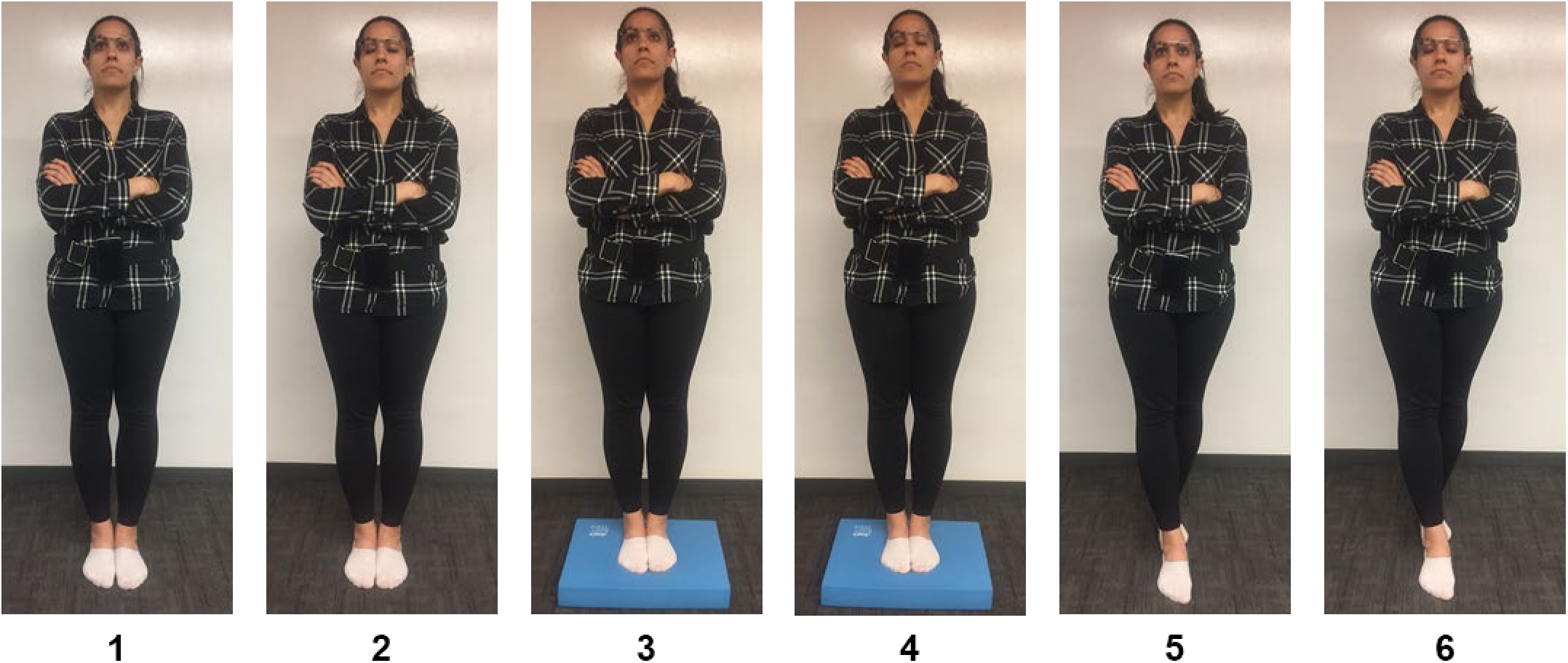
BAM protocol conditions. (1, 2) Feet together, standing on a firm surface used for conditions 1 (eyes open) and 2 (eyes closed). (3, 4) Feet together, standing on a foam surface used for conditions 3 (eyes open) and 4 (eyes closed). (5, 6) Feet in tandem stance, standing on a firm surface used for conditions 5 (eyes open) and 6 (eyes closed).

### Data acquisition

An Android application was developed to synchronize recording of device IMU data between smartglasses and the waist-mounted smartphone. The application was loaded on both Google Glass and the Android smartphones prior to testing. The application allowed Google Glass to pair with a smartphone via Bluetooth. Messages sent via Bluetooth from Google Glass to the smartphone were used to initiate a timer on Google Glass and begin storing IMU accelerometer and gyroscope values (sampled at 50 Hz). When running on Google Glass, the application provides instructions on stance, a timer, and a tone that plays at the end of each stance session.

### Data analysis

The first 10 seconds of data were discarded to ensure stability of measures (50 seconds of data total). Accelerometer data (ACC) from each trial was low-pass filtered using a phase less 4th order, Butterworth filter with a cut-off frequency of 1.25 Hz [13]. The normalized path length (NPL; mG/s; higher values indicate more sway) of the acceleration time series from the anterior-posterior (AP) postural sway data was defined as:

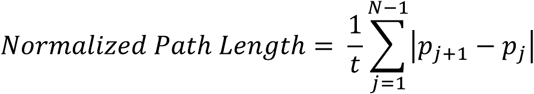

where *t* is the time duration, *N* is the number of time samples, and *p_j_* is the ACC at time sample *j* in the AP direction (z-axis on both devices, **Fig 2**). NPL was also calculated from the combined ACC magnitude.

**Fig. 2.**
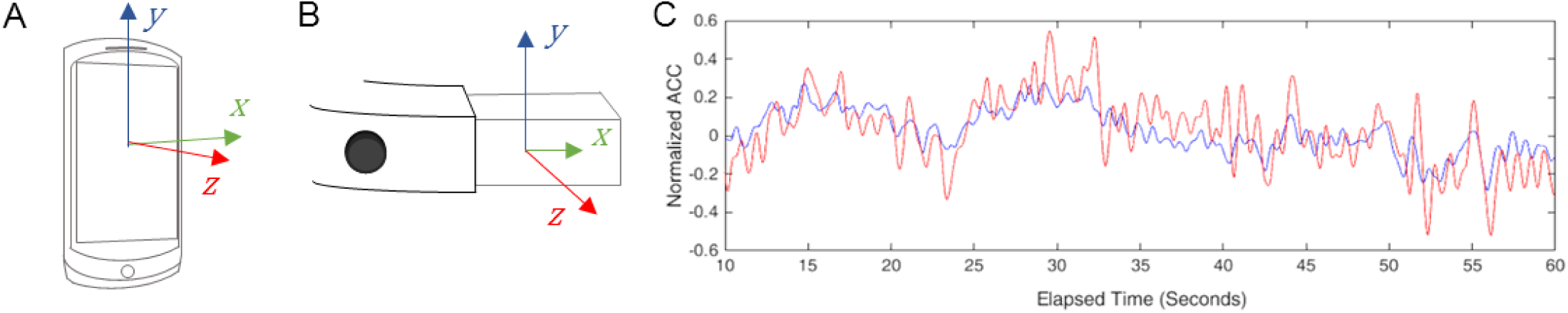
ACC collection with smartphone and smartglasses. A. Axes of accelerometer on Android smartphone compared with (B) Google Glass. C. Example comparison of low-pass filtered ACC (z-axis) collected during a trial of condition 6 in Glass (red) compared with waist-mounted smartphone (blue).

Smartglasses-based measurements of NPL along different axes were compared with smartphone measurements using Spearman’s rank correlation coefficient [15]. For comparison of differences between stances, the nonparametric Kruskal-Wallis test was used to compare mean ranks [16, 17]. Normality of measurements within stance conditions was evaluated by the Anderson-Darling test [18]. Significant differences between correlation coefficients were determined by treating them as Pearson coefficients and using the standard Fisher’s z-transformation to compare using a standard normal procedure [19]. Test-retest reliability of NPL measurements was estimated for each condition between the two sessions by calculating the intraclass correlation coefficient (ICC) and corresponding 95% confidence intervals (CI) [20, 21]. NPL was standardized as previously described [13]:

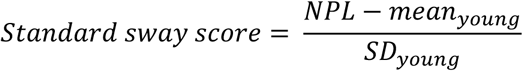

where *mean_young_* is the mean of young healthy subjects (age 18-34 years) on condition 1 (reported previously as 9.4 mG/sec), and *SD_young_* is the corresponding standard deviation (2.2). The composite score, also as previously described, was defined as the sum of the standard sway scores across all six conditions.

## Results

### Smartglasses-based measurement of AP sway correlates with waist-based measurement

All 42 subjects completed two balance trials on conditions 1 through 3, like previous reports [13]. Both trials of the eyes closed/foam surface condition (condition 4) were successfully completed by 37 subjects (88%). One subject failed a trial of the eyes open/tandem stance condition (condition 5). 30 subjects (71%) completed two trials of the eyes closed/tandem stance condition (condition 6). Overall, two trials on all 6 conditions were completed by 28 (66%) subjects.

NPL AP sway measured from the head was strongly correlated (Spearman’s rank correlation coefficient = 0.85) with NPL AP sway measured from the waist (**Fig 3A**). Mean NPL AP sway measured from the waist (**Fig 3B**) was in good agreement with previously reported values [13], although we observed a higher mean for condition 6. The mean (SD) composite score was 21.4 (18.0), which was in good agreement with the previously reported value of 19.6 (15.3) for healthy subjects.

**Figure 3:**
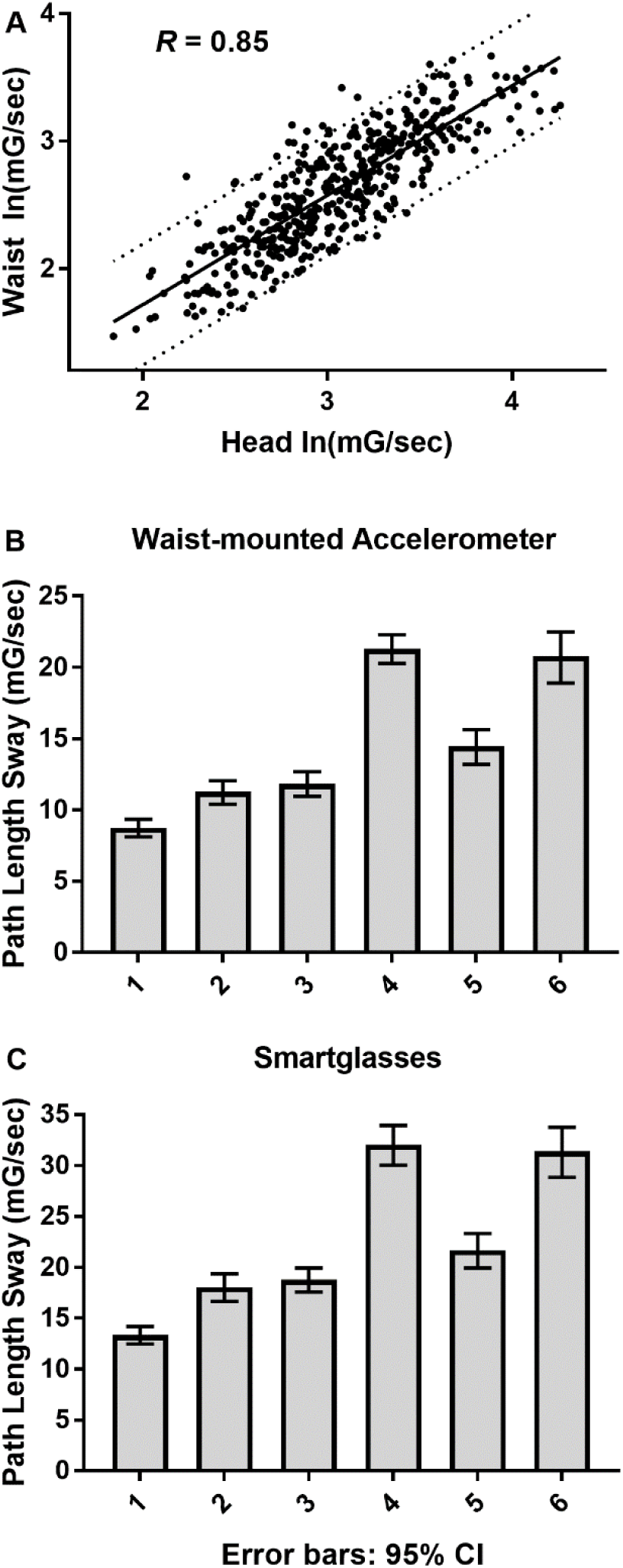
AP sway. A. AP sway measured from the head was strongly correlated with AP measured from the waist (pooled data from all conditions with 95% prediction bands). Geometric mean and 95% CI for waist-based (B) and head-based (C) measurement of AP NPL by condition.

While NPL measured from the head was generally larger than NPL measured from the waist in each trial, mean NPL AP sway measured from the head in each condition (**Fig 3C**) was observed to follow a similar trend as the means measured from the waist. Signification differences (Kruskal-Wallis, α = 0.05) were found between each set of eyes open and eyes closed conditions, as well as between standing on feet together/firm surface compared with foam surface or tandem stance.

### Correlation between head and waist measurements was significantly stronger when comparing the NPL calculated from the combined magnitude of all three ACC axes

Measuring sway along the ACC’s AP axis was previously shown to be sufficient to differentiate healthy subjects from subjects with vestibular disorders [13]. However, the additional ACC acquired from commercial off-the-shelf smart devices may further enhance measurement accuracy, particularly along the mediolateral x-axis. Indeed, the NPL calculated using all three axes (total NPL) was found to have a significantly stronger correlation (Spearman’s rank correlation coefficient = 0.90) between head- and waist-based measurements (**Fig 4A**). Mean total NPL measured in each condition followed similar trends as using AP NPL only for both waist- (**Fig 4B**) and head-based (**Fig 4C**) measurements.

**Figure 4:**
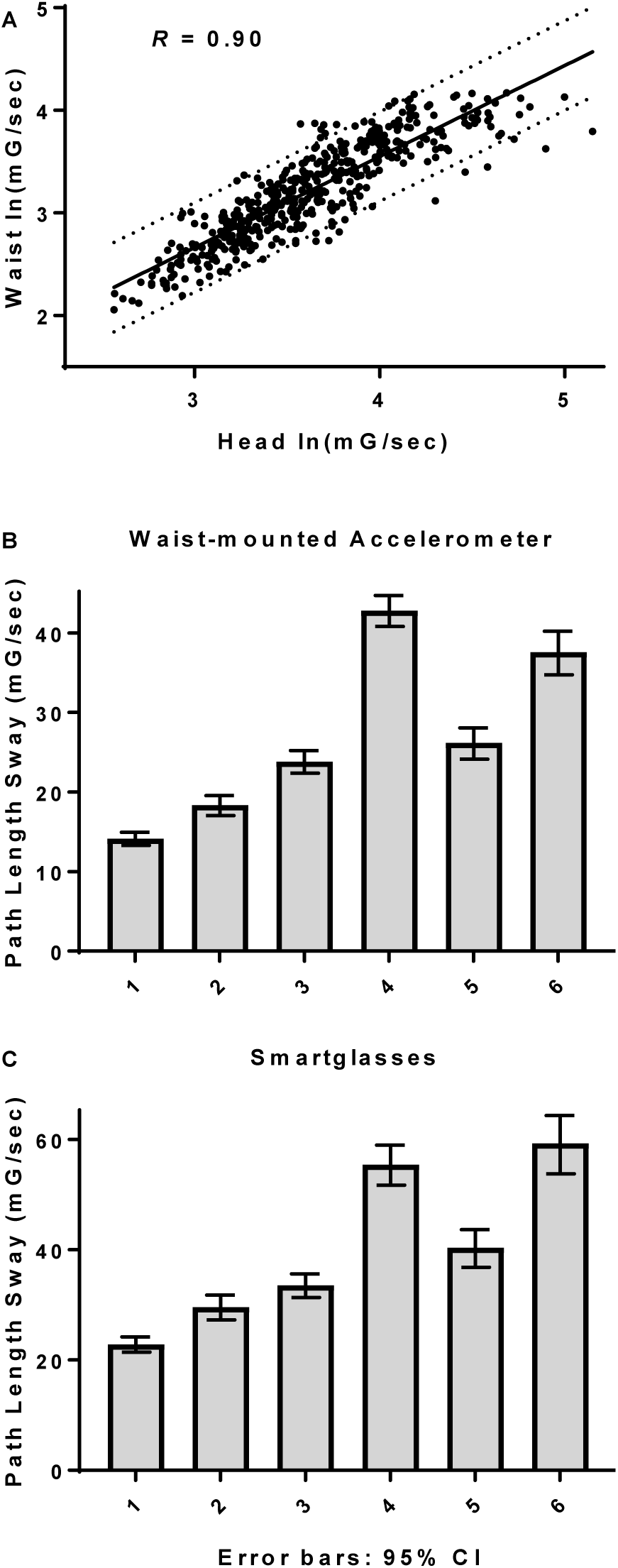
Total NPL magnitude. A. Total sway measured from head was more strongly correlated with sway measured from the waist. Geometric mean and 95% CI for waist-based (B) and head-based (C) measurement of total NPL by condition.

### Test-retest reliability of measures of NPL sway were comparable between head and waist

Previously, the same day test-retest reliability of NPL AP measured from the waist was found to be generally good (ICC ≥ 0.74) across all conditions except for condition 6 [13]. Here, same day test-retest reliability of AP NPL measured from the head with smartglasses (**Fig 5A**) was found to be very good, with an ICC (95% CI) of 0.85 (0.81-0.88). This was comparable to our estimation of the test-retest reliability of waist-based AP NPL (**Fig 5B**), which was 0.84 (0.80-0.88), agreeing with previously reported values. Using total NPL, we found a slight improvement in test-retest reliability in both head (**Fig 5C**) and waist-based measurements (**Fig 5D**). ICC (95%) was found to be 0.87 (0.83-0.90) in the case of head-based measurement, as opposed to 0.90 (0.88-0.92) in the case of waist-based measurement.

**Figure 5:**
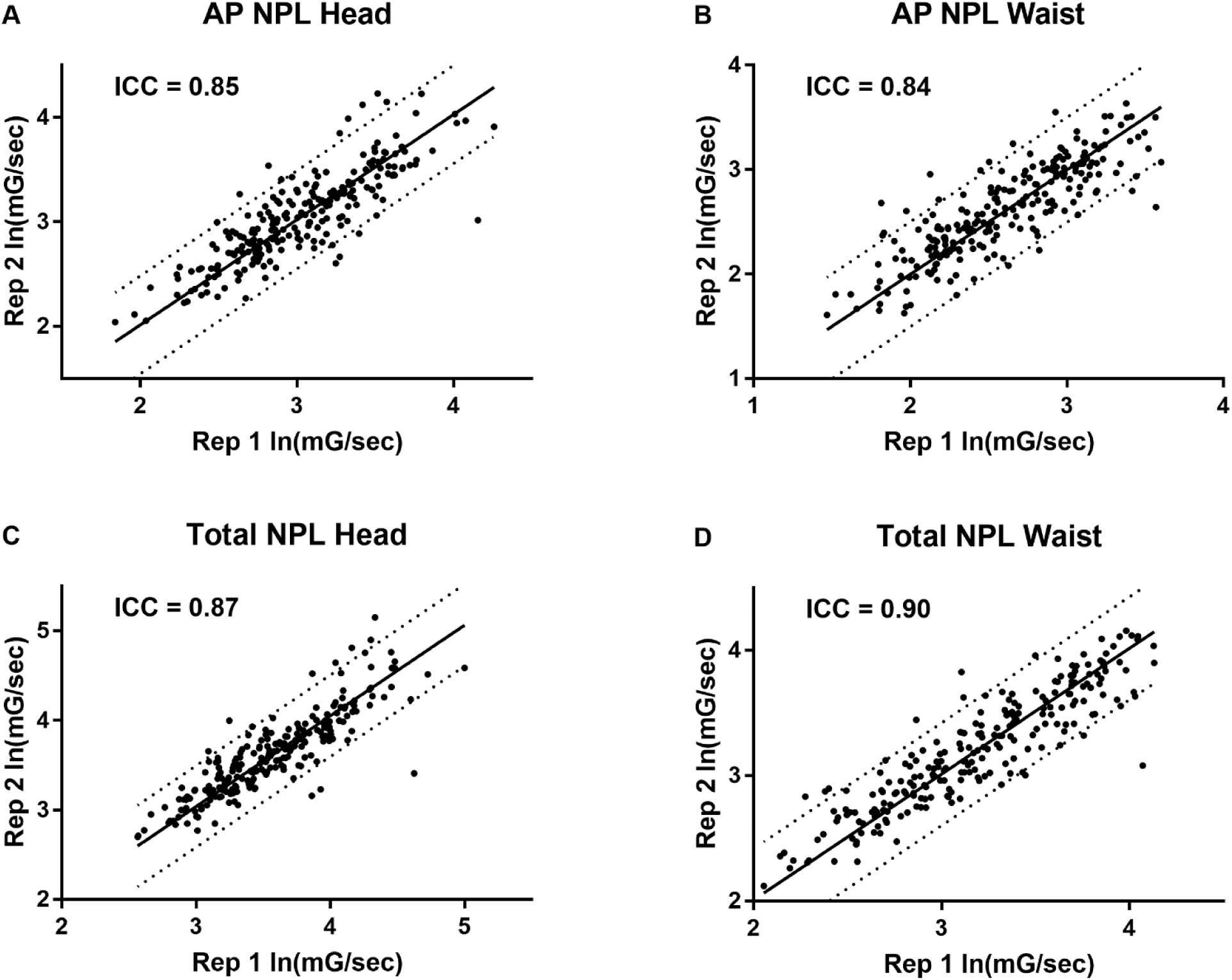
Head-based BAM has comparable test-retest reliability to waist-based BAM. Test-retest ICC for accelerometer measures of postural sway along AP (A = head, B = waist) and all axes (C = head, D = waist).

## Discussion

This study indicates that head-based measurement of AP or total sway following the BAM protocol produces similar results to waist-based measurement. This included similar relative differences between test conditions as well as similar test-retest reliability. All subjects successfully complete the study, demonstrating the feasibility of using a head-mounted wearable to assess balance, at least in a healthy control population.

Condition effects previously supported the validity of waist AP NPL as a measure of balance. Measured sway was larger with eyes closed versus eyes open for all stance conditions. Sway was also larger in tandem stance and on foam surface conditions compared with corresponding conditions with feet together on a firm surface. These condition effects indicated that ACC measured from the waist was sensitive to changes in the sensory modalities available for balance, including vision and somatosensation [22]. Postural sway as measured by waist-based BAM was adequate to discriminate between persons with peripheral vestibular impairments from those without balance-impairment [13]. While the specific stance conditions used in the BAM protocol were not found to discriminate between healthy and concussed adolescents, instrumenting the Balance Error Scoring System (BESS) protocol with a waist-based inertial sensor led to superior diagnostic classification of recently concussed individuals compared with BESS alone [23].

Head-based measurement of balance using smartglasses offers a promising opportunity to develop a balance assessment tool that has integrated instruction and feedback. Smartglasses include not only Google Glass, but an evolving spectrum of devices from both major device consumer device manufacturers, including Epson (Moverio) and Intel (Recon Jet), as well as companies dedicated to the smartglasses market, including Vuzix and ODG. Because of their form-factor, smartglasses could offer vestibular rehabilitation through gamified virtual reality (VR) better than other head-mounted systems. In general, VR is a promising technology for treatment of balance disorders, including vestibular deficits associated with concussion/TBI [24-31]; however, non-transparent VR headsets completely block external visual stimuli and are typically bulky, which could limit their utility in vestibular therapy. In contrast, Google Glass weighs only 1.3 ounces –ten times less than the consumer-grade VR Oculus Rift. Thus, demonstrating that smartglasses are capable of objectively assessing balance deficits could lead to an integrated system for vestibular assessment and rehabilitation. In conclusion, further studies are warranted to demonstrate smartglasses ability to distinguish balance disorders, including those stemming from concussion.

## Acknowledgments

Funded in part by Department of Defense/Defense Health Program SBIR Phase II contract #W81XWH-14-C-0007 (awarded to TIAX, LLC).

